# A Tandem Fluorescent Rh1::eGFP–mCherry Reporter Reveals Rhodopsin 1 Trafficking and Turnover In Vivo in Drosophila Photoreceptors

**DOI:** 10.64898/2026.10.09.757923

**Authors:** Hanieh Gheflati, Christine Steiner, Nilofar Feizy

**Affiliations:** University of Hohenheim, Institute of Biology, Department of Biochemistry, Ottilie-Zeller-Weg 2, 70599 Stuttgart, Germany; Lead contact: Nilofar Feizy, University of Hohenheim, Institute of Biology, Ottilie-Zeller-Weg 2, 70599 Stuttgart, Germany

**Keywords:** *Drosophila* photoreceptors, protein degradation, Photoreceptor homeostasis lysosomal pathway, Rh1

## Abstract

Rhodopsin 1 (Rh1), the major visual pigment of *Drosophila* photoreceptors, requires a vitamin A-derived chromophore for proper folding, maturation and transport to the rhabdomere. While the endocytic trafficking of mature, light-activated Rh1 to endolysosomal compartments is well characterized, the fate of immature Rh1 during defective ER maturation remains unclear. Here, we developed a Rh1::eGFP-mCherry tandem reporter to monitor Rh1 localization and compartment acidification in vivo. Exploiting the acid stability of mCherry, this reporter enables visualization of Rh1 after delivery to acidic environments and complements conventional antibody-based approaches. Under physiological conditions, pH-dependent quenching of eGFP fluorescence distinguished rhabdomeric Rh1 from Rh1 in acidic endolysosomal compartments. In vitamin A-deficient flies, immature apo-Rh1 accumulated within Calnexin-positive ER membranes, where we identified discrete acidified subdomains characterized by loss of eGFP fluorescence while mCherry fluorescence persisted. These structures remained continuous with the ER but were distinct from conventional endolysosomal compartments, revealing previously unrecognized acidified ER domains associated with immature Rh1. Together, our findings identify acidified ER subdomains as a previously unrecognized stage in Rh1 quality control and establish Rh1::eGFP-mCherry as a robust in vivo reporter for investigating rhodopsin trafficking, maturation and turnover under physiological and disease-relevant conditions.

## INTRODUCTION

Rhodopsin 1 (Rh1), the major visual pigment of *Drosophila melanogaster* outer photoreceptors, is synthesized in the endoplasmic reticulum (ER), trafficked through the secretory pathway and delivered to the rhabdomere, a specialized apical membrane compartment required for phototransduction (O’Tousa et al., 1985; Zuker et al., 1985). Proper synthesis, folding, trafficking and turnover of Rh1 are essential for photoreceptor homeostasis, whereas defects in any of these processes lead to Rh1 mislocalization, cellular stress and progressive retinal degeneration (Alloway et al., 2000; Chinchore et al., 2009; Satoh and Ready, 2005).

A critical step in Rh1 biogenesis is the covalent incorporation of the vitamin A-derived chromophore 11-cis-3-hydroxyretinal. Chromophore binding promotes Rh1 folding, maturation and stability, enabling efficient export from the ER and delivery to the rhabdomere. Accordingly, vitamin A deficiency causes defective Rh1 maturation and dramatically reduces rhabdomeric Rh1 levels (Huber et al., 1994; Ozaki et al., 1993; Colley et al., 1995). Although mature, light-activated Rh1 undergoes well-characterized endocytic trafficking to endolysosomal compartments, the intracellular fate of immature apo-Rh1 during impaired chromophore-dependent maturation remains poorly understood.

Most current knowledge of Rh1 localization and trafficking has been obtained using antibody-based immunolocalization and genetic analysis of trafficking mutants (Alloway et al., 2000; Chinchore et al., 2009; Iwanami et al., 2016).

While these methods have provided important mechanistic insights, they do not readily allow visualization of Rh1 as it moves between intracellular compartments in living photoreceptors or enters acidic environments during trafficking and degradation. Consequently, it remains unclear whether immature Rh1 is exclusively retained for ER-associated degradation or whether additional trafficking pathways contribute to its processing under chromophore-deficient conditions.

Tandem fluorescent reporters combining a pH-sensitive GFP variant with an acid-stable red fluorescent protein provide a powerful strategy to monitor protein delivery into acidic compartments. Originally developed to measure autophagic flux (Kimura et al., 2007), this approach has subsequently been applied to selected membrane cargo proteins in cultured cells (Parenti et al., 2017), but its application to physiological membrane trafficking in vivo has remained limited.

Here, we generated the Rh1::eGFP–mCherry tandem reporter to visualize Rh1 localization and compartment acidification simultaneously in living *Drosophila* photoreceptors. Using this reporter, we investigated Rh1 trafficking under physiological conditions and during vitamin A deprivation. Our findings reveal previously unrecognized acidified ER subdomains associated with immature Rh1 and establish Rh1::eGFP–mCherry as a versatile in vivo tool for studying GPCR trafficking, maturation and degradation.

## RESULTS AND DISCUSSION

### Validation of Rh1::eGFP–mCherry as an *in vivo* reporter for rhodopsin trafficking and localization

To generate an *in vivo* reporter for monitoring Rh1 trafficking, a DNA construct was generated encoding Rhodopsin 1 (Rh1) fused at its C-terminus to the tandem fluorescent proteins eGFP and mCherry. Expression of the fusion protein was driven by the endogenous *rhodopsin 1 (ninaE)* promoter in R1–R6 photoreceptor cells of *Drosophila* (Fig. 1A). Transgenic fly lines expressing Rh1::eGFP::mCherry were established either in a wild-type background containing endogenous Rh1 or in a *ninaE^17^* mutant background lacking native Rh1.

**Fig. 1.**
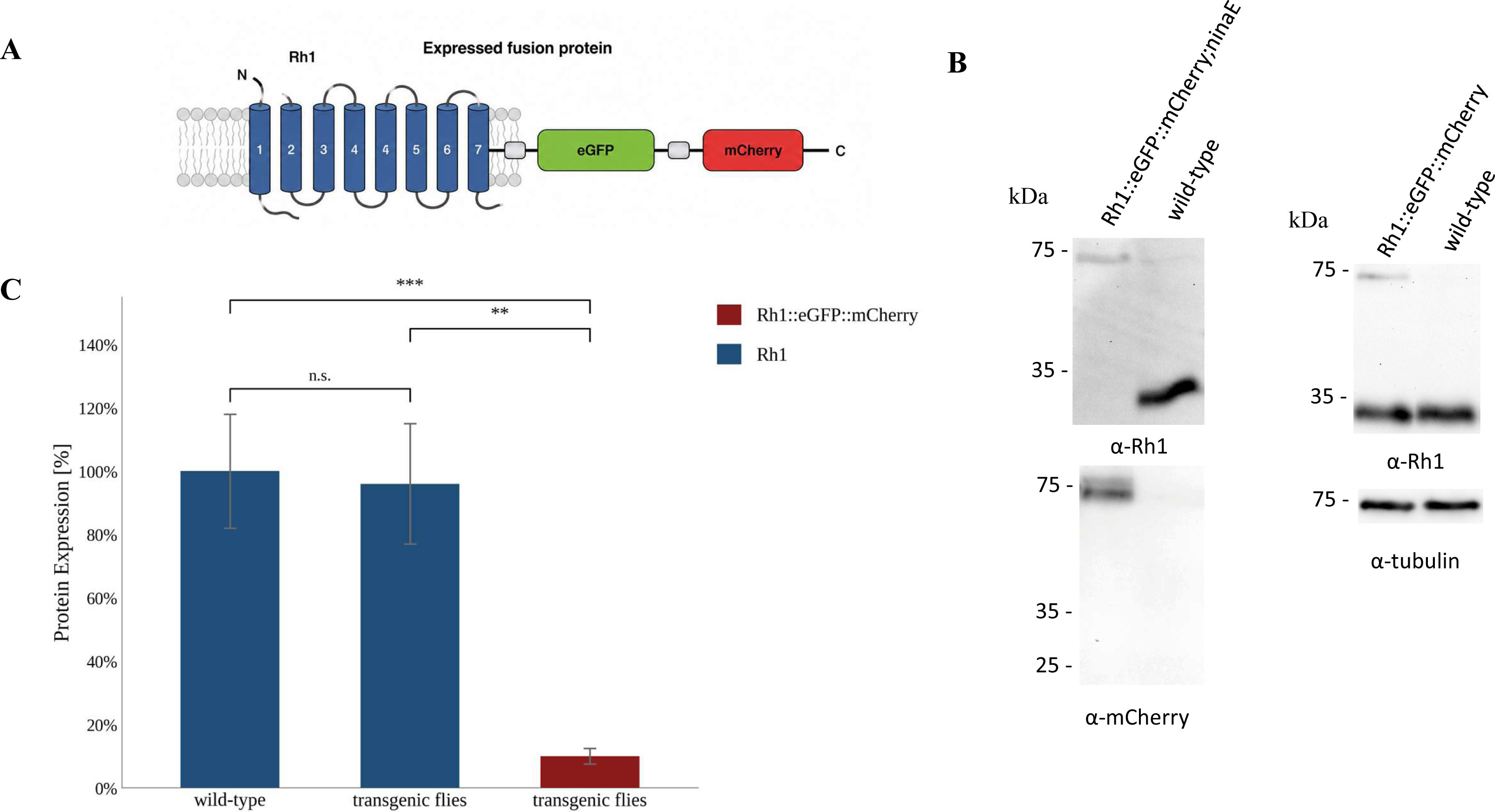
Generation of Rh1::eGFP::mCherry-expressing transgenic *Drosophila*. **(A)** Schematic representation of the C-terminal Rh1::eGFP::mCherry fusion protein. A DNA construct was generated encoding Rhodopsin 1 (Rh1) fused at its C-terminus to tandem fluorescent proteins eGFP and mCherry. **(B)** Immunoblot analysis assessing expression of the Rh1::eGFP::mCherry fusion protein in heads of transgenic flies. Rh1 and Rh1::eGFP::mCherry were detected using a polyclonal Rh1 antibody (α-Rh1). Expression of Rh1::eGFP::mCherry was additionally detected using an α-mCherry antibody. **(C)** Quantification of endogenous Rh1 and Rh1::eGFP::mCherry protein levels in wild-type flies compared with flies expressing Rh1::eGFP::mCherry in a wild-type background. Immunoblot analysis was performed using α-Rh1. Rh1 levels in wild-type flies were normalized to 100%. Data represent mean values from three independent biological replicates (n = 3; ns, not significant; **P < 0.002; ***P < 0.0002).

To confirm expression of the Rh1::eGFP::mCherry fusion protein in photoreceptor cells, immunoblot analyses were performed using protein extracts prepared from adult fly heads. A polyclonal rabbit anti-Rh1 antibody was used to detect both endogenous Rh1 and fusion protein, whereas a mouse anti-mCherry antibody specifically detected the reporter fusion. Rh1::eGFP::mCherry was detected as a distinct band at the expected molecular weight of approximately 75 kDa (Fig. 1B). Quantification of immunoblot signals revealed that the fusion protein was expressed at approximately 10% of the endogenous Rh1 level (Fig. 1C). Comparable expression levels were observed in the presence or absence of endogenous Rh1, indicating that transgene expression was not substantially influenced by the endogenous *ninaE* locus.

### Functional assessment of Rh1::eGFP::mCherry by electroretinography

To assess whether the Rh1::eGFP::mCherry fusion protein retains functional properties of Rh1, electroretinography (ERG) recordings were performed. Following dark adaptation, flies were sequentially stimulated with a single 5 s orange light pulse followed by the orange–blue–blue– orange–orange (OBBOO) light protocol (Fig. 2A; see Materials and Methods).

**Fig. 2.**
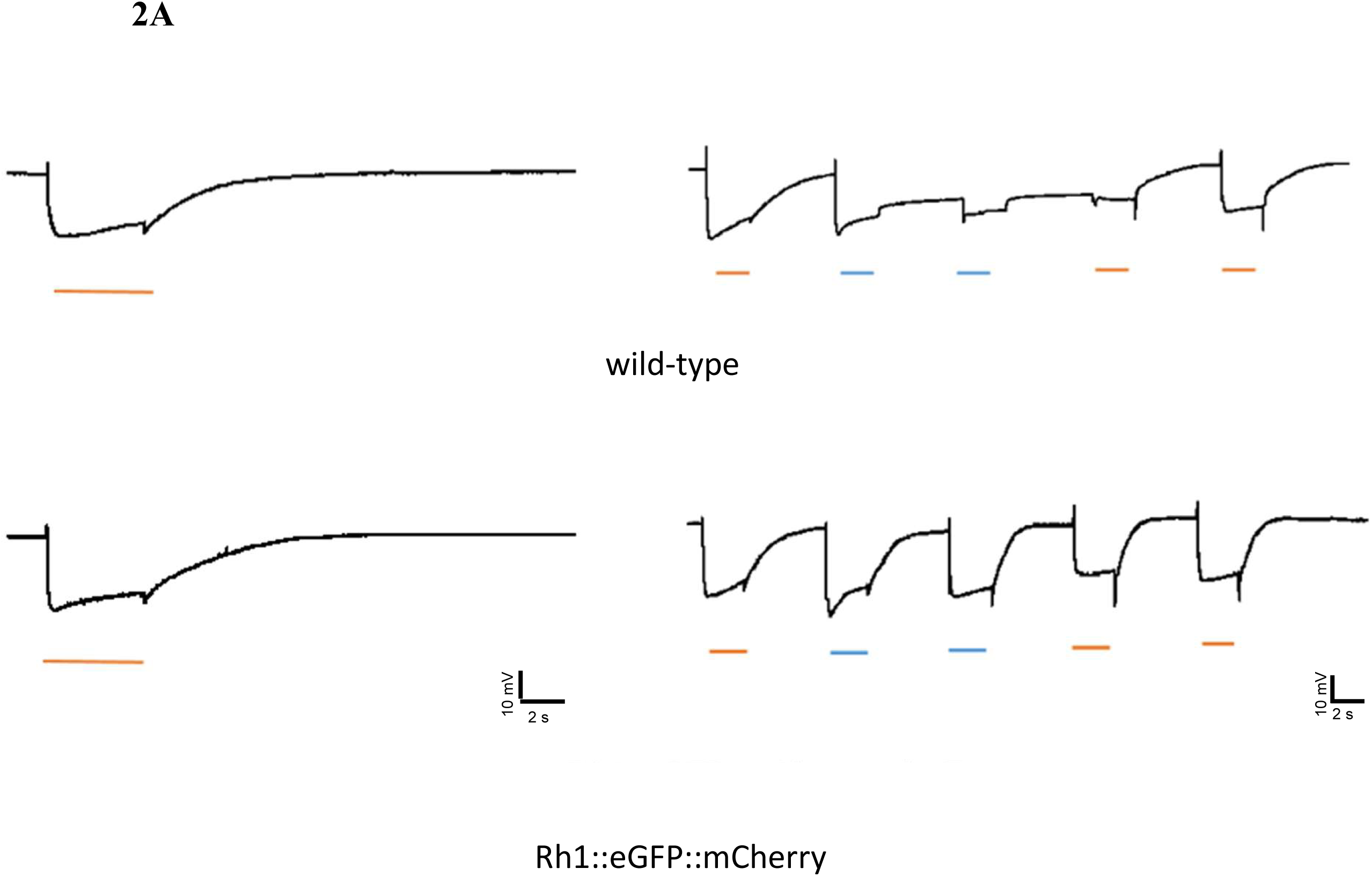
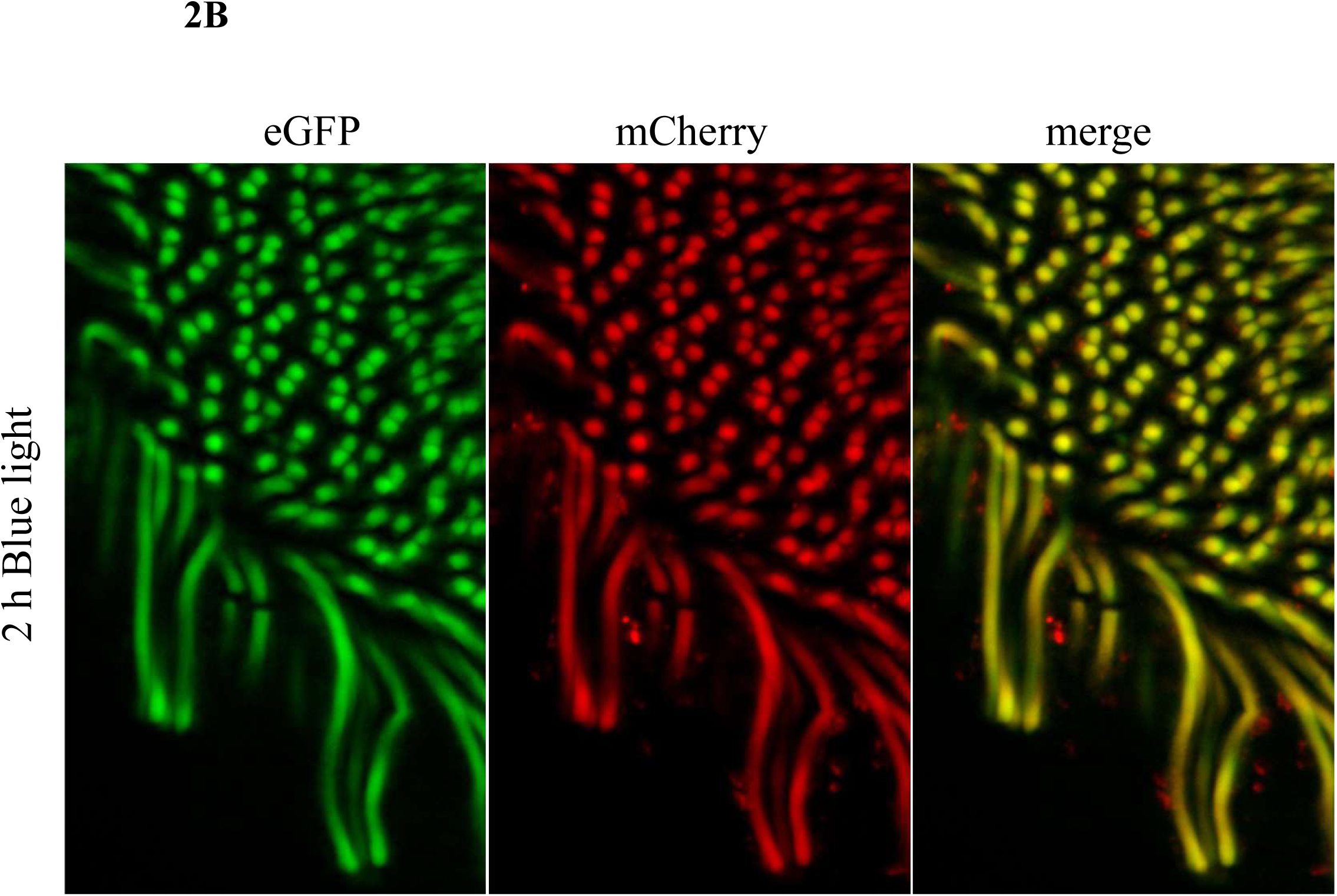
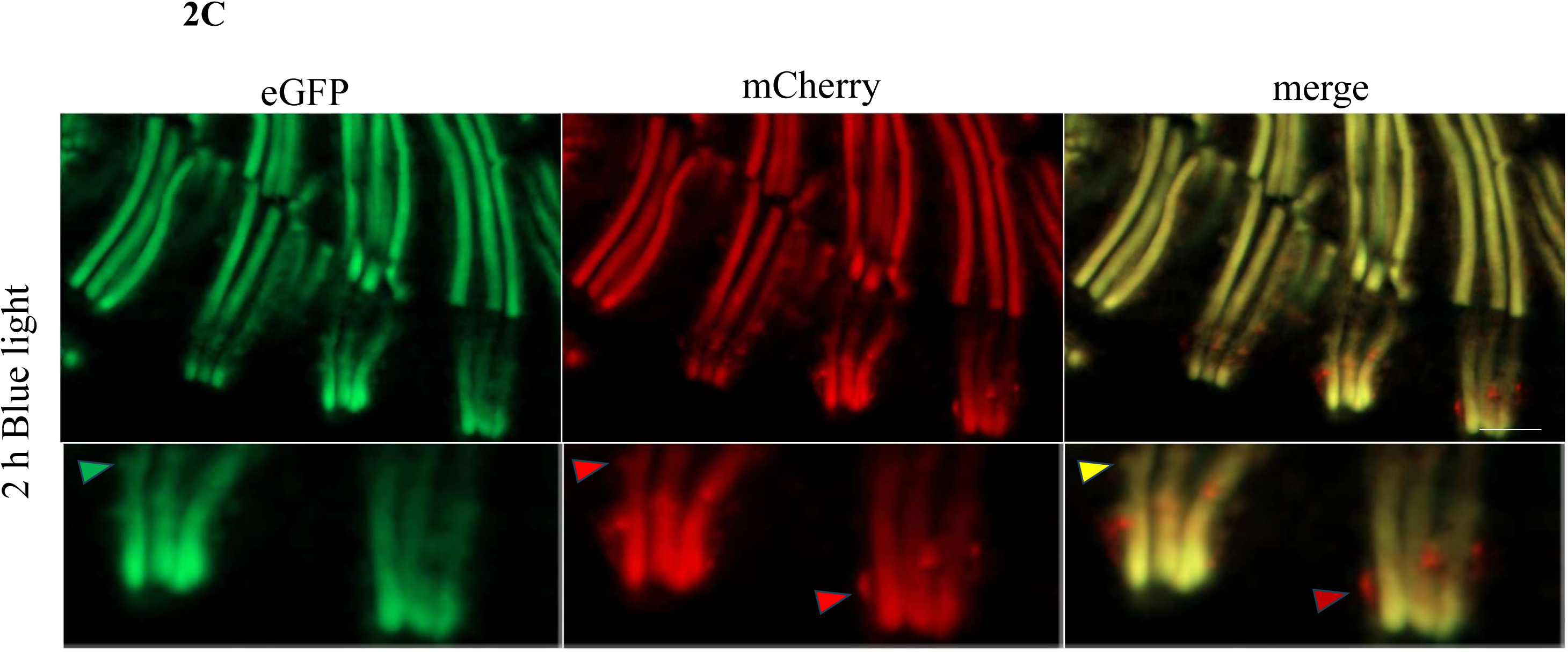
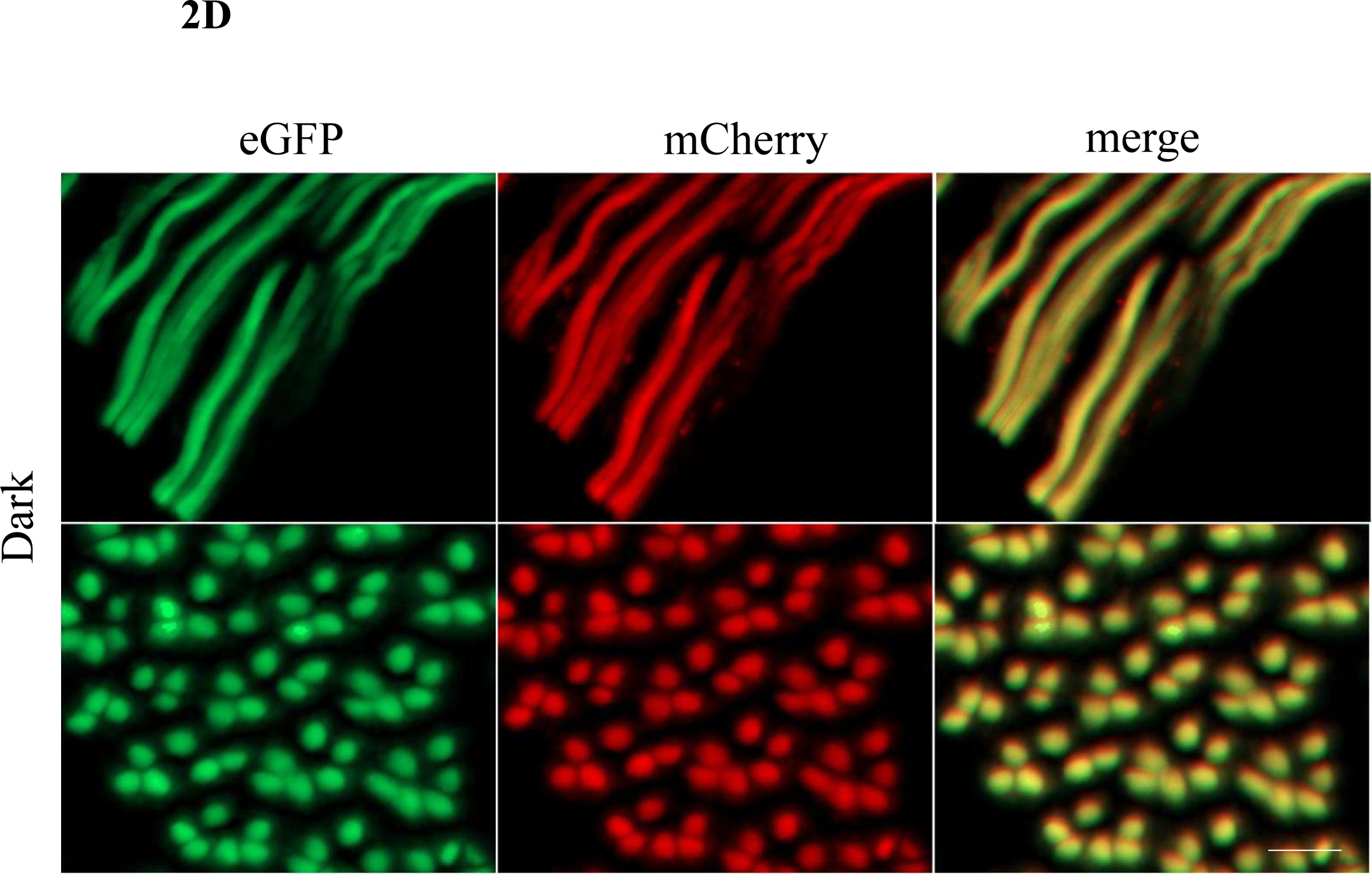
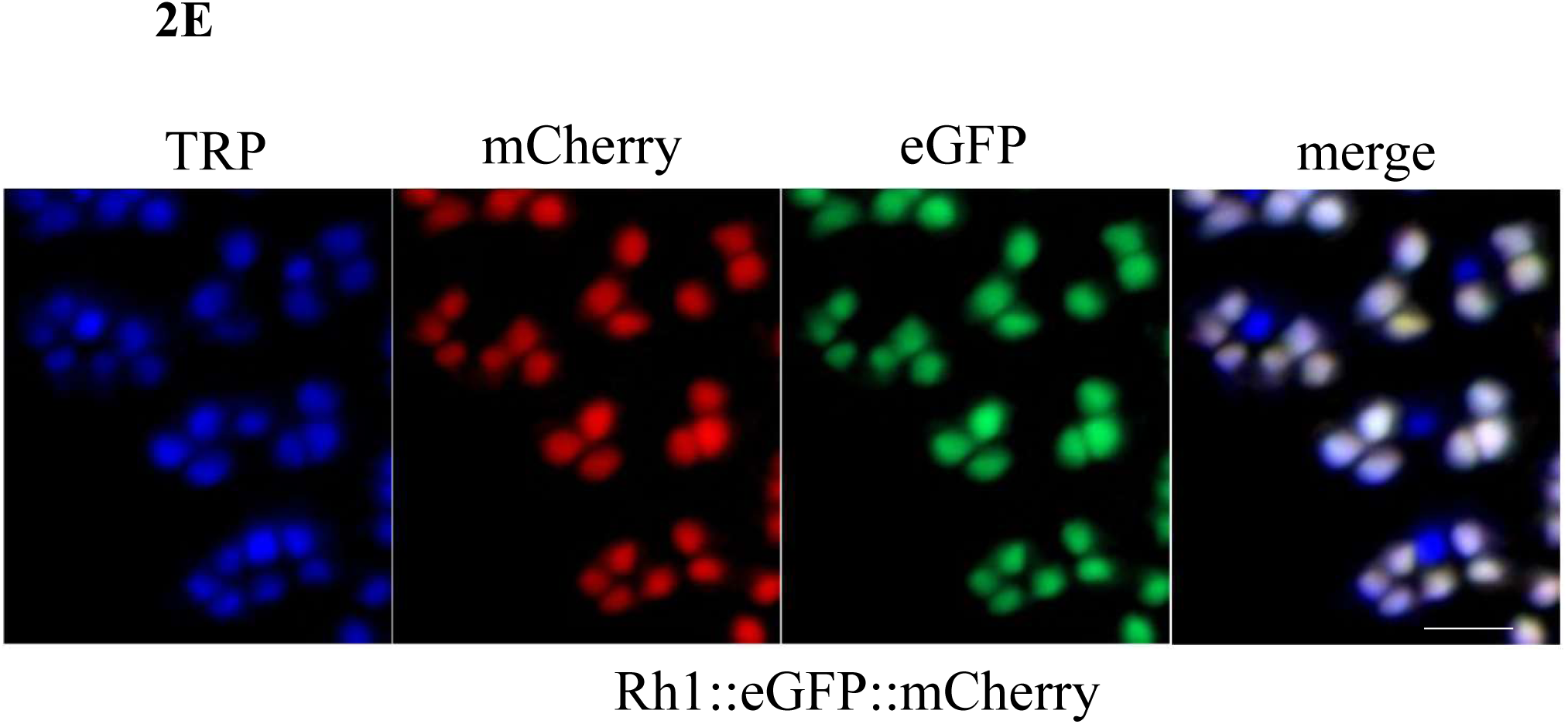
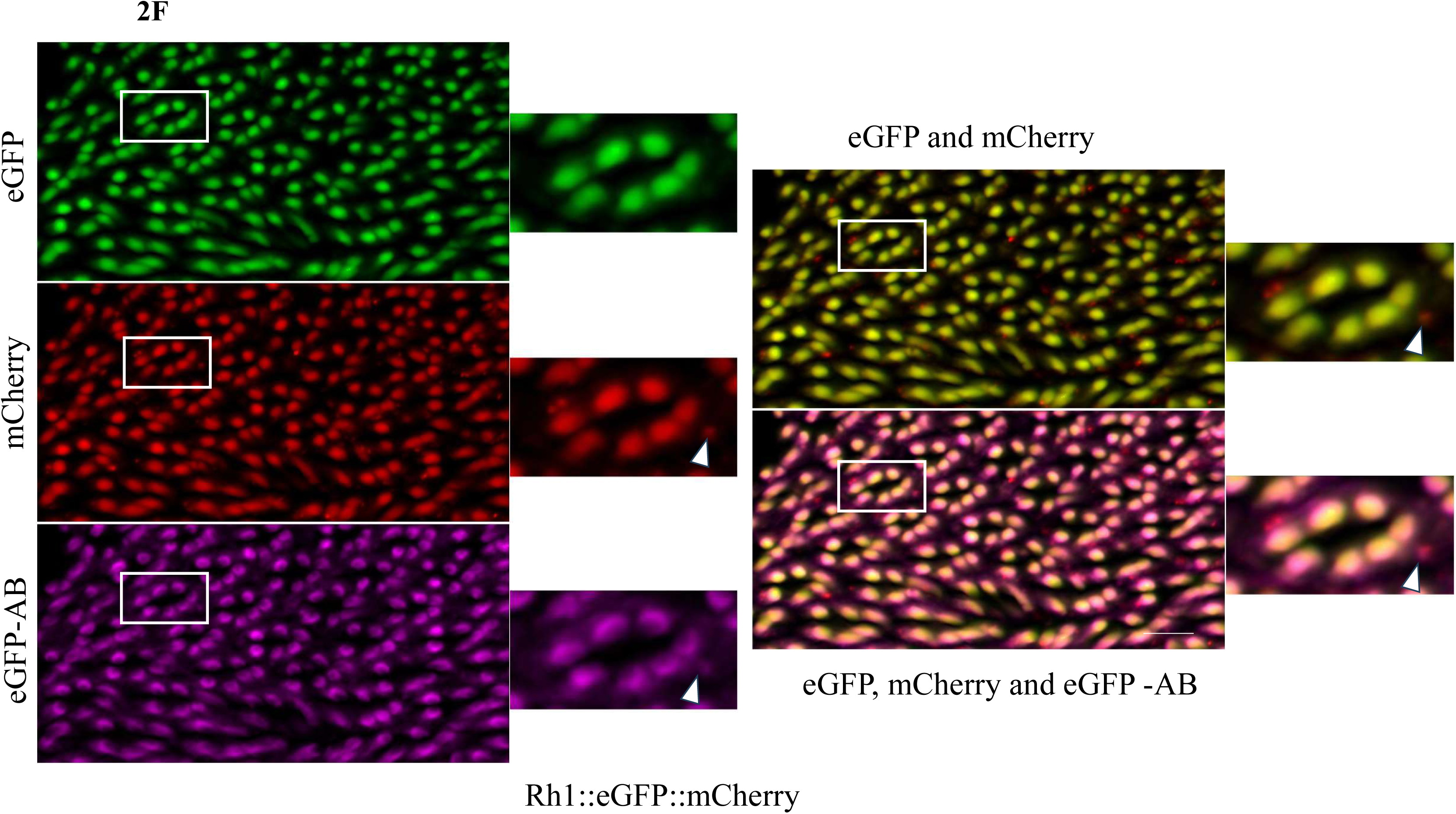
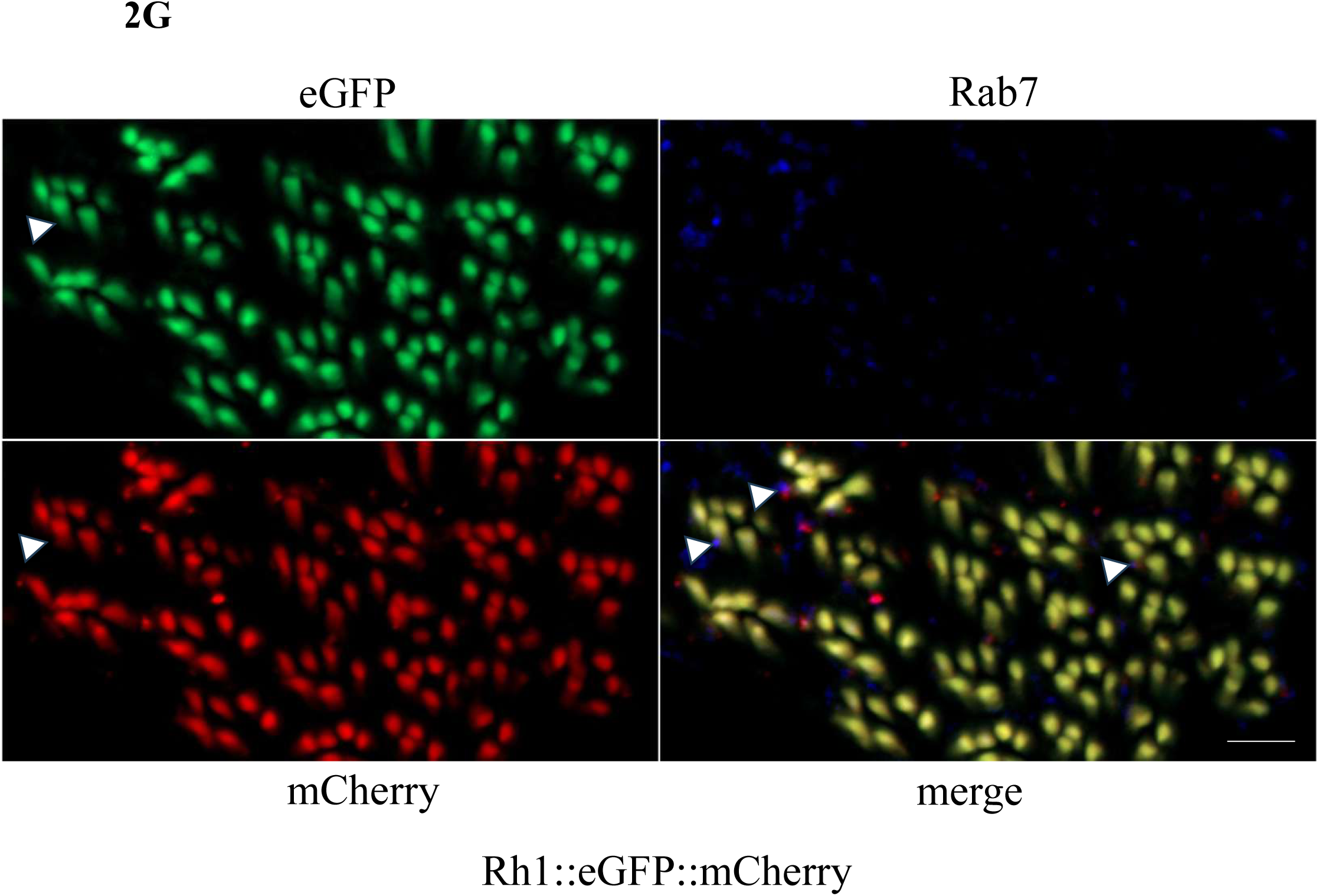
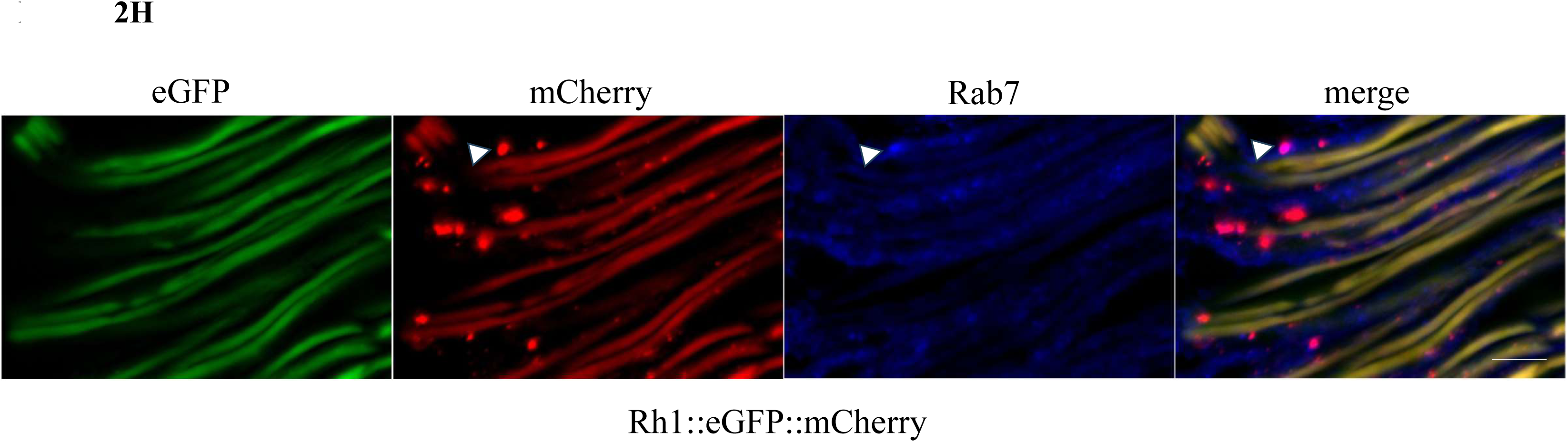
Validation of Rh1::eGFP–mCherry as an *in vivo* reporter for Rh1 localization, trafficking, and delivery to acidic compartments. (A) Electroretinograms (ERG) of 3-day-old wild-type and transgenic (Rh1::eGFP::mCherry;ninaE) flies. Dark-adapted flies were subjected to a single orange light stimulus (left side) or to an OBBOO protocol. Orange and blue light stimuli are indicated by orange and blue bars, respectively. Flies expressing Rh1::eGFP::mCherry in the absence of endogenous Rh1 (Rh1::eGFP::mCherry;ninaE) displayed wild-type-like light responses, with no prolonged depolarization afterpotential (PDA) observed following blue light stimulation. (B-D). Rh1::eGFP–mCherry reports Rh1 trafficking into acidic intracellular compartments. Representative retinal sections of photoreceptors expressing Rh1::eGFP::mCherry following 2 h of blue light exposure (B-C) or under dark-adapted conditions (D). Native eGFP fluorescence (green), mCherry fluorescence (red), and merged images are shown. In dark-adapted photoreceptors, cytoplasmic mCherry-positive vesicles were readily detected. Compared with dark-adapted conditions, light-stimulated photoreceptors contained a substantially higher number of intracellular vesicles displaying strong mCherry fluorescence (red arrowhead) accompanied by markedly reduced or absent eGFP signal. A subset of intracellular vesicles retained both eGFP and mCherry fluorescence (yellow arrowhead), consistent with non-acidified or incompletely acidified trafficking intermediates. The green arrowhead indicates an intracellular vesicle that is detectable by native eGFP fluorescence, representing a non-acidified Rh1-containing compartment. (E) Rh1::eGFP–mCherry is correctly targeted to the rhabdomeres. Representative retinal section of a dark-adapted photoreceptor expressing Rh1::eGFP::mCherry. TRP immunostaining (blue) marks the rhabdomeres, while native mCherry (green) and eGFP (red) fluorescence label the Rh1 fusion protein. The complete co-localization of Rh1::eGFP::mCherry with TRP demonstrates correct rhabdomeral targeting of the fusion protein, and the complete overlap of eGFP and mCherry fluorescence indicates that both fluorophores remain associated under physiological conditions. (F)Anti-GFP immunostaining demonstrates retention of the GFP moiety in acidic Rh1-containing compartments. Representative retinal section of a light-stimulated photoreceptor expressing Rh1::eGFP::mCherry. Native eGFP fluorescence (green), mCherry fluorescence (red), and anti-GFP immunostaining (magenta) are shown individually and in merged images. Many intracellular mCherry-positive vesicles lacking native eGFP fluorescence remained positive for anti-GFP immunostaining, demonstrating that the GFP moiety is retained in acidic compartments and that the loss of native eGFP fluorescence primarily reflects pH-dependent quenching. In contrast, a subset of mCherry-positive vesicles was negative for both native eGFP fluorescence and anti-GFP immunostaining, suggesting degradation of the GFP moiety or masking/loss of the GFP epitope in more advanced degradative compartments. Boxed regions indicate representative intracellular vesicles. **(G-H)** Rab7 labels a subset of acidic Rh1-containing late endosomes. Representative retinal section of a light-stimulated photoreceptor expressing Rh1::eGFP::mCherry and immunostained for Rab7. Native eGFP fluorescence (green), mCherry fluorescence (red), Rab7 immunostaining (blue), and the merged image are shown. Arrowheads indicate mCherry-positive, eGFP fluorescence-negative vesicles that co-localize with Rab7, identifying them as late endosomes. A subset of mCherry-positive vesicles lacked Rab7 immunoreactivity, suggesting that Rh1 is also present in additional degradative compartments. Scale bars, 10 µm in all panels.

Upon orange light stimulation, photoreceptor cells of all genotypes exhibited rapid depolarization that was maintained throughout the duration of the stimulus, indicating intact phototransduction responses. In wild-type flies, blue light illumination induced a pronounced prolonged depolarization afterpotential (PDA), reflecting efficient photoconversion of Rh1 to metarhodopsin and accumulation of active metarhodopsin that is not immediately quenched by Arrestin 2.

In contrast, flies expressing Rh1::eGFP::mCherry in the *ninaE^17^* mutant background lacking endogenous Rh1 did not display a detectable PDA following blue light stimulation. This absence of PDA is consistent with the substantially lower expression level of the fusion protein compared with endogenous Rh1 and suggests that the amount of Rh1::eGFP::mCherry expressed from the transgene is insufficient to generate enough metarhodopsin to generate a PDA.

### Rh1::eGFP::mCherry is correctly targeted to the rhabdomere and reports Rh1 trafficking into acidic compartments

The subcellular localization of Rh1::eGFP::mCherry was examined by fluorescence microscopy (Fig. 2B–E). Rh1::eGFP::mCherry was predominantly localized to the rhabdomeres of R1–R6 photoreceptor cells, as confirmed by co-staining with the rhabdomeric marker TRP (Fig. 2E). The complete overlap of eGFP and mCherry fluorescence within the rhabdomeres demonstrates correct targeting of the fusion protein and suggests that the fluorescent tags do not interfere with Rh1 localization.

In dark-adapted photoreceptors, cytoplasmic mCherry-positive vesicles were readily detected, indicating basal intracellular Rh1 trafficking under physiological conditions (Fig. 2D). Compared with dark-adapted photoreceptors, light-stimulated photoreceptors contained a substantially higher number of intracellular vesicles displaying strong mCherry fluorescence accompanied by markedly reduced or absent eGFP signal (Fig. 2B–C). These vesicles also appeared noticeably larger after light stimulation, consistent with increased endolysosomal activity accompanying enhanced Rh1 internalization and degradation. This fluorescence pattern is consistent with increased delivery of Rh1 into acidic compartments following light-induced internalization (Kiselev et al., 2000; Han et al., 2007; Xiong and Bellen, 2013), where eGFP fluorescence is quenched while the mCherry signal remains largely preserved. A subset of vesicles retained both eGFP and mCherry fluorescence (Fig. 2C, yellow arrowheads), consistent with early endosomal intermediates that had not yet undergone complete acidification.

To determine whether the loss of eGFP fluorescence in Rh1::eGFP::mCherry-positive vesicles resulted from pH-dependent quenching or degradation of the GFP moiety, retinal sections were immunostained using an anti-GFP antibody. Although many intracellular mCherry-positive vesicles lacked detectable native eGFP fluorescence, these same vesicles remained readily detectable by anti-GFP immunostaining (Fig. 2F, arrowhead). This demonstrates that the GFP component remains associated with the fusion protein in the majority of Rh1::eGFP::mCherry-positive vesicles following delivery to intracellular compartments and that the loss of native eGFP fluorescence primarily reflects pH-dependent quenching rather than degradation.

However, a subset of mCherry-positive vesicles was detected by neither native eGFP fluorescence nor anti-GFP immunostaining. These vesicles may represent more advanced degradative compartments in which either the GFP moiety or the GFP epitope has undergone proteolytic degradation, while the more acid-resistant mCherry signal remains detectable. Thus, the Rh1::eGFP::mCherry reporter enables discrimination between Rh1-containing compartments at different stages of maturation and degradation and provides a dynamic readout of Rh1 trafficking from the rhabdomere to acidic intracellular compartments.

To further characterize these intracellular compartments, retinal sections were immunostained for the late endosomal marker Rab7. A subset of mCherry-positive vesicles lacking detectable native eGFP fluorescence co-localized with Rab7, identifying these structures as late endosomal compartments (Fig. 2G–H, arrowheads). However, not all mCherry-positive vesicles were Rab7-positive, suggesting that a proportion of Rh1-containing vesicles represents other intracellular compartments, including more mature lysosomal compartments (Satoh & Ready, 2005; Wang et al., 2014).

It should be noted that immunostaining experiments, such as those performed for Rab7 and GFP, require fixation, detergent-mediated membrane permeabilization, and antibody incubation. These procedures may alter the luminal pH of intracellular compartments and could potentially restore native eGFP fluorescence that had previously been quenched under acidic conditions. Consequently, immunostained preparations are only of limited suitability for assessing pH-dependent fluorescence changes. This potential limitation does not affect the interpretation of the Rab7 co-localization analysis. The purpose of the Rab7 experiments was to identify the mCherry-positive vesicles lacking detectable native eGFP fluorescence, demonstrating that a subset of these compartments corresponds to late endosomes. Since not all mCherry-positive vesicles were Rab7-positive, additional Rh1-containing compartments are likely present, including more mature lysosomal compartments that were not identified by Rab7 immunostaining. These observations are consistent with previous studies showing that light-activated Rh1 traffics through late endosomes before lysosomal degradation, while a smaller fraction is recycled to the rhabdomere via retromer-dependent trafficking (Wang et al., 2014).

Thus, direct imaging of native eGFP and mCherry fluorescence in unfixed retinal tissue represents the most reliable approach for monitoring compartment acidification, whereas immunohistochemistry primarily serves to define the molecular identity of Rh1-containing compartments.

### Vitamin A deficiency reveals reversible acidified ER-associated Rh1 domains

To investigate the influence of chromophore availability on Rh1 maturation and intracellular organization, Rh1::eGFP–mCherry-expressing fly larvae were raised under vitamin A-deficient conditions and were maintained on the same diet after eclosion for one additional day before analysis. Under these conditions, Rh1::eGFP–mCherry remained predominantly associated with Calnexin-positive membranes of the endoplasmic reticulum (ER), consistent with previous reports that chromophore-free Rh1 apoprotein accumulates as an immature, ER-retained species in the absence of vitamin A (Huber et al., 1994; Ozaki et al., 1993; Colley et al., 1995). (Fig. 3A, left). Most of the reticular ER network displayed overlapping eGFP and mCherry fluorescence, indicating retention of the intact fusion protein within the ER.

**Fig. 3.**
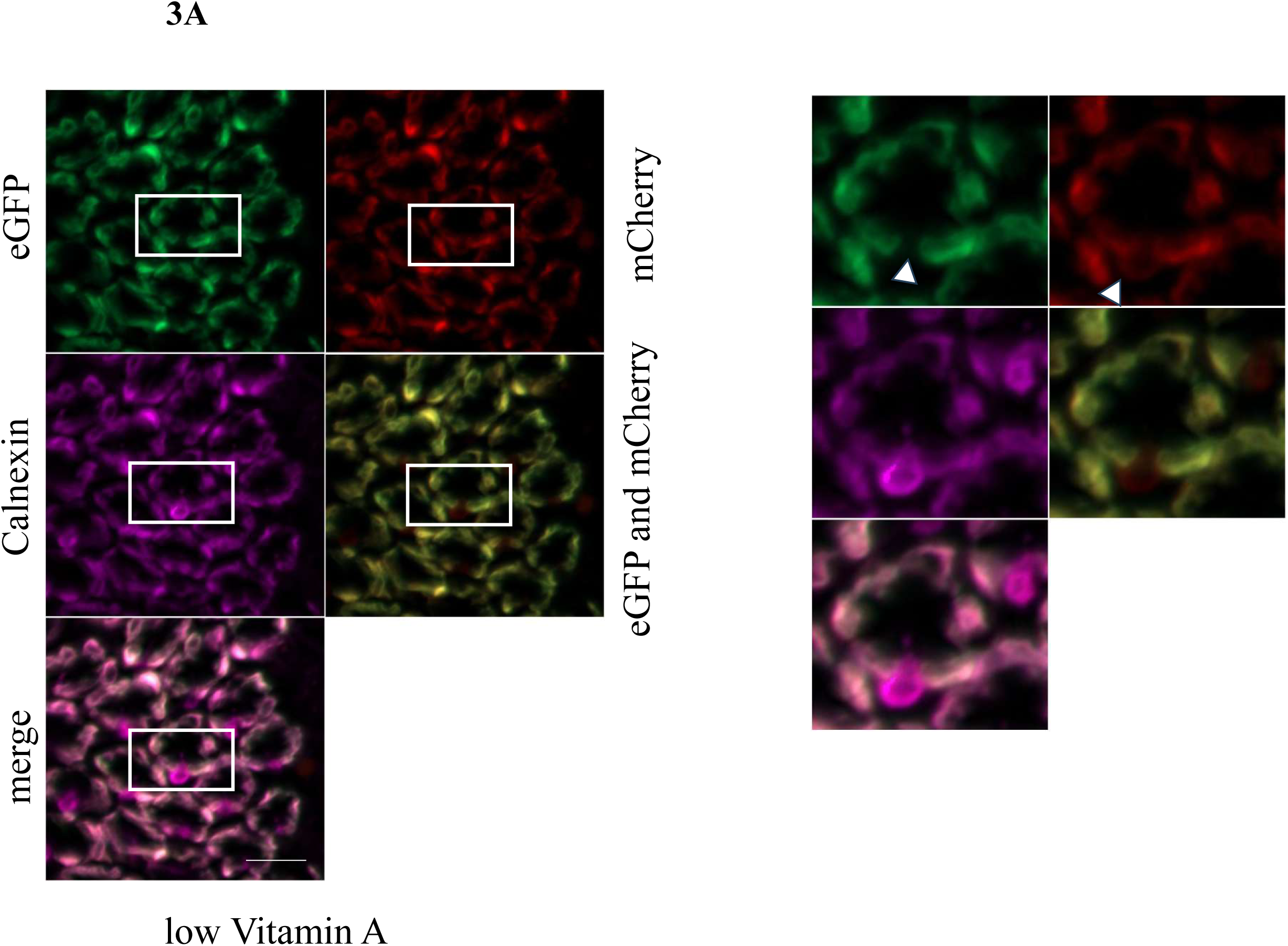
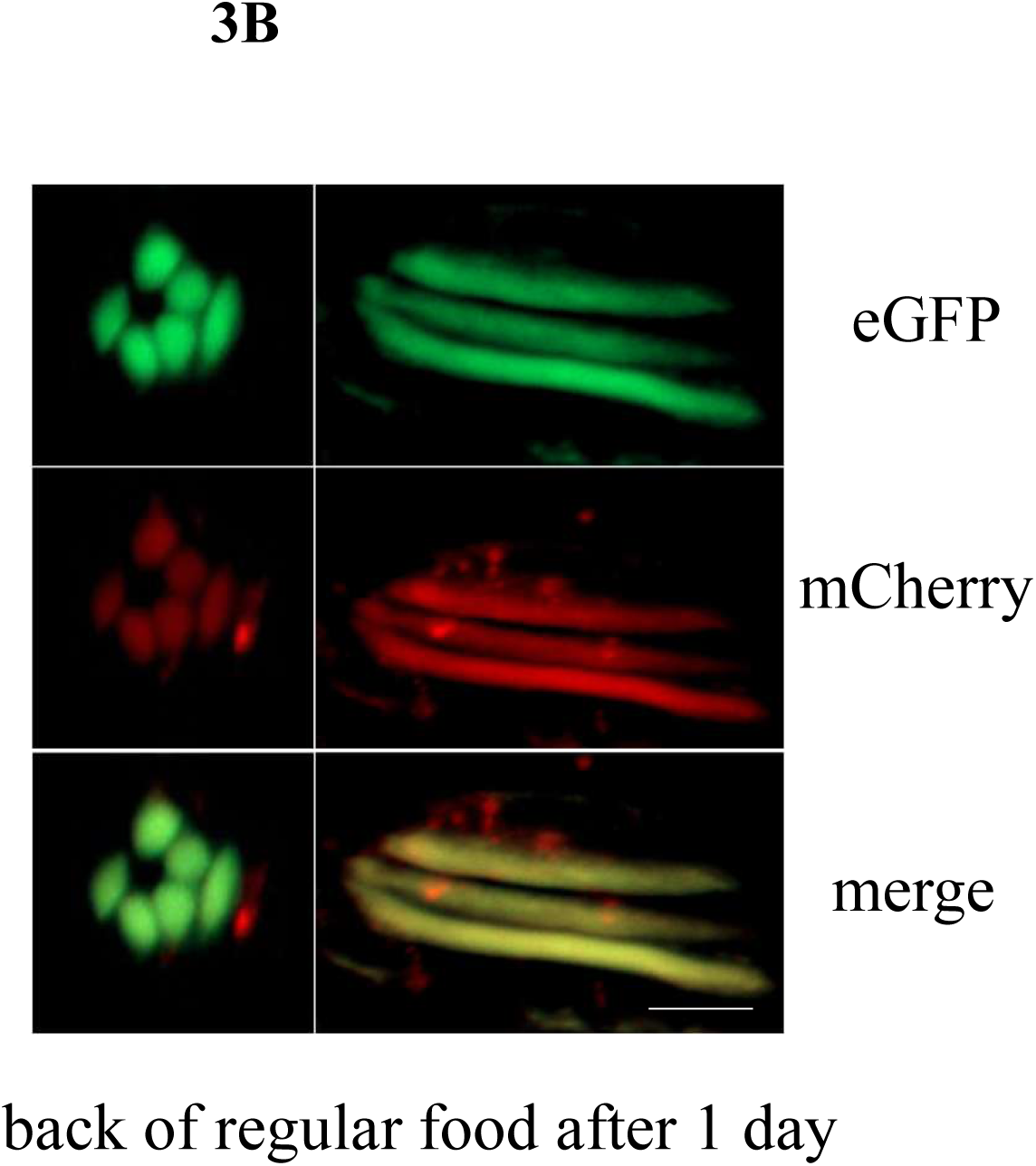
Vitamin A deficiency reveals acidified Rh1-containing ER-associated domains. **(A)** Representative confocal images of photoreceptor cells expressing Rh1::eGFP–mCh**e**rry under vitamin A-deficient conditions. Panels show native eGFP fluorescence (green), native mCherry fluorescence (red), the merged eGFP/mCherry signals, Calnexin immunostaining (magenta) labeling the endoplasmic reticulum (ER), and the final merged image. Rh1::eGFP– mCherry is predominantly retained within Calnexin-positive ER membranes, consistent with accumulation of immature apo-Rh1. While most of the ER exhibits overlapping eGFP and mCherry fluorescence, distinct Calnexin-positive ER regions display a reduced eGFP fluorescence, indicative of localized acidification causing pH-dependent quenching of eGFP. The panels on the right show magnified views of the boxed regions outlined in the left panels. **(B)**Vitamin A supplementation restores Rh1 maturation and normal intracellular trafficking.Representative confocal images of photoreceptor cells expressing Rh1::eGFP– mCherry following recovery from vitamin A deficiency. Vitamin A-deficient flies were transferred to standard food for two days before imaging. Rh1::eGFP–mCherry localization to the rhabdomere was restored, indicating recovery of Rh1 maturation and proper rhabdomeral targeting. Scale bars, 10 µm

Interestingly, distinct Calnexin-positive regions exhibited locally reduced eGFP fluorescence while maintaining the continuous reticular morphology of the ER (Fig. 3A, right). In contrast to the small, discrete mCherry-positive vesicles observed following light-induced Rh1 internalization under physiological conditions, these structures were larger, irregularly shaped, and remained integrated within the ER network rather than appearing as independent vesicular compartments. Their strong co-localization with Calnexin indicates that these acidified Rh1-containing domains remain part of, or are closely associated with, the ER rather than representing conventional endolysosomal vesicles.

Our validation experiments (Fig. 2F-H) demonstrated that the loss of native eGFP fluorescence primarily reflects pH-dependent quenching rather than degradation of the reporter. Therefore, the reduced eGFP/mCherry fluorescence ratio observed within these ER-associated domains is consistent with localized acidification. Because the ER lumen is generally considered to maintain a near-neutral pH, these observations suggest the existence of localized acidic microenvironments within specific Rh1-containing regions of the ER and reveal a previously unrecognized functional heterogeneity of the ER under conditions of impaired Rh1 maturation.

The molecular identity of these acidified ER-associated domains remains unknown. One possible explanation is that they represent specialized ER quality-control compartments that transiently sequester immature Rh1. In light of the findings of Zhao et al. (2023), who demonstrated that chronic ER stress can induce IRE1/XBP1-dependent selective ER-phagy, it is conceivable that these locally acidified Rh1-containing domains correspond to early intermediates of ER quality-control pathways or membrane domains associated with ER-phagy. However, because neither autophagic nor ER-phagy markers were examined in the present study, our data do not directly demonstrate activation of these pathways. The relationship between these acidified ER domains and ER-phagy therefore remains speculative and will require future investigation using established molecular markers.

To determine whether the accumulation of acidified ER-associated Rh1 domains under vitamin A deficiency represents a reversible adaptive state rather than an irreversible consequence of Rh1 misfolding, flies were kept on a vitamin A-deficient diet for one day after eclosion and subsequently transferred to standard food for two days before imaging. Strikingly, this recovery period was sufficient to restore Rh1::eGFP–mCherry localization to the rhabdomere. Moreover, following light stimulation, numerous acidic mCherry-positive vesicles were again observed, demonstrating re-establishment of the normal light-dependent endolysosomal trafficking pathway (Fig. 3B). These findings indicate that ER retention of apo-Rh1 is reversible and that chromophore re-supplementation is sufficient to restore Rh1 maturation, rhabdomeral targeting, and subsequent light-induced trafficking. Particularly relevant to our findings are studies that established the Blue Light-Induced Chromophore Supply (BLICS) approach. In this approach, flies are initially maintained under carotenoid-deficient conditions, resulting in the accumulation of Rh1 apoprotein in the ER in the absence of chromophore. Subsequent supplementation with all-trans-retinal and its blue light-induced isomerization to 11-cis-retinal allows Rh1 maturation to be triggered synchronously, enabling the subsequent transport of Rh1 from the ER through the Golgi to the rhabdomere to be followed (Satoh et al., 1997; Iwanami et al., 2016).

Unlike conventional ER stress reporters, which primarily report activation of stress signaling pathways, Rh1::eGFP–mCherry directly visualizes the intracellular fate of Rh1 while simultaneously reporting changes in the luminal environment of Rh1-containing compartments. This genetically encoded reporter therefore enables the visualization of Rh1 maturation, ER retention, light-dependent endolysosomal trafficking, and compartment acidification within a single experimental system. Furthermore, the identification of localized acidified ER-associated Rh1 domains reveals a previously unrecognized aspect of Rh1 quality control and highlights the potential of this reporter to uncover cellular mechanisms of receptor maturation and intracellular quality control that are not readily accessible using conventional UPR or ER stress reporters alone.

## CONCLUSION

The present study establishes Rh1::eGFP–mCherry as a versatile in vivo reporter for simultaneously monitoring Rh1 localization, trafficking, and the luminal environment of Rh1-containing compartments in *Drosophila* photoreceptors. Unlike conventional ER stress reporters, which provide indirect information about cellular stress pathways, this tandem fluorescent reporter directly visualizes the subcellular localization and luminal pH environment of Rh1-containing compartments throughout its maturation and trafficking cycle.

Under vitamin A deficiency, the absence of chromophore resulted in the accumulation of immature apo-Rh1 within the endoplasmic reticulum. In addition to classical ER retention, we identified previously undescribed ER-associated acidified Rh1 domains characterized by localized loss of eGFP fluorescence while retaining mCherry signal. Their persistent association with Calnexin-positive ER membranes suggests that these structures represent specialized regions of the ER with altered luminal properties rather than conventional endolysosomal compartments.

The rapid disappearance of these domains following vitamin A restoration, together with the recovery of rhabdomeric Rh1 localization and light-dependent endolysosomal trafficking, demonstrates that impaired Rh1 maturation under chromophore deficiency is a highly reversible process. These findings indicate that photoreceptor cells can dynamically adapt to limitations in Rh1 maturation by reorganizing Rh1-containing ER compartments.

Although the molecular mechanisms underlying the formation and function of these acidified ER-associated Rh1 domains remain to be determined, their identification provides new insight into ER quality control and Rh1 homeostasis. Future studies investigating their relationship with ER-phagy, autophagy, and other quality control pathways will be essential to define their biological role.

Overall, this work reveals an unexpected layer of complexity in Rh1 maturation and ER organization and highlights the importance of directly monitoring protein localization and compartmental properties to understand mechanisms maintaining photoreceptor proteostasis.

## MATERIALS AND METHODS

### Fly husbandry and light conditions

Adult flies were reared on either a standard yeast–cornmeal diet or a low-vitamin A diet, both supplemented with methylparaben, vitamin C, and propionic acid. The standard diet consisted of 84 g/L fresh baker’s yeast, 50 g/L cornmeal, 32 g/L sucrose, and 12 g/L agar. The low-vitamin A diet was prepared by dissolving 10 g dry yeast, 10 g sucrose, 2 g agar, and 20 mg cholesterol in 100 mL of double-distilled water. The suspension was heated in a microwave with intermittent stirring until all components were completely dissolved and subsequently cooled to approximately 60 °C. After cooling, 64 μL of 10% methylparaben (Nipagin) dissolved in ethanol, 500 μL propionic acid, and a small amount of ascorbic acid (approximately one spatula tip) were added. All fly stocks were reared and maintained under standard laboratory conditions at 25 °C.

The fly strains employed in this study included wild-type (w), the Rh1 mutant ninaE17 (y w; ninaE17), yw;;Rh1::eGFP-mCherry, yw;Rh1::eGFP and Rh1::eGFP-mCherry;ninaE.

For dark adaptation, flies were maintained at 25 °C in complete darkness for the specified durations. Blue light adaptation was achieved by exposing flies to blue light (310–490 nm; Ee₄₇₀ nm = 28 μW/cm²) for 2h inside a transparent colored plastic chamber illuminated with a white fluorescent lamp (4000 K, 1750 lux; Ee470 nm = 298 μW/cm², Ee590 nm = 215 μW/cm²; Osram, Munich, Germany).

Dark-adapted flies, as well as flies illuminated with red light, were manipulated under dim red illumination provided by a cold light source (KL 1500 LCD, Schott, Mainz, Germany) fitted with a deep-red long-pass filter (RG630, Schott). This setup was used to avoid unintended photoconversion of Rh1. In contrast, light-adapted flies were processed under standard ambient white laboratory lighting.

### Cloning and Expression of Rh1::eGFP::mCherry

Transgenic flies expressing the Rh1::eGFP::mCherry fusion protein were generated using a previously established Rh1-based expression construct as the template (Feizy et al., 2024). Based on this parental construct, the final Rh1::eGFP::mCherry expression cassette was assembled through three sequential cloning steps.

Initially, the Rh1 cDNA was inserted into the expression vector. In the second cloning step, the eGFP coding sequence lacking a stop codon was fused in-frame to the C-terminus of Rh1, allowing continuous translation of the fusion protein. Finally, the mCherry coding sequence was cloned downstream of eGFP and terminated with a stop codon, yielding the complete Rh1::eGFP::mCherry fusion construct.Cloning was performed using the restriction enzymes *BamHI, EcoRI, XhoI,* and *XbaI*.

Correct assembly of the plasmid was confirmed by DNA sequencing. Subsequently, the Rh1::eGFP::mCherry construct, initially cloned into the pENTR™1A vector, was transferred into a modified YC4 destination vector by site-specific recombination between att sites using the Gateway® cloning system (Invitrogen, Germany). The modified YC4 vector contained the *Drosophila* Rh1 promoter (base pairs −833 to +67), attR recombination sites, and the terminal 0.6 kb of the Rh1 3′ untranslated region (Richter et al., 2011). P-element-mediated germline transformation of the *Drosophila melanogaster* yellow white (yw) host strain was performed by the Drosophila Injection Service at FlyORF (Zurich, Switzerland).

### Protein Extraction from Fly Heads

For protein isolation, ten fly heads were homogenized in 40 μL of SDS extraction buffer containing 4% SDS, 1 mM EDTA, and 75 mM Tris-HCl (pH 6.8). The homogenates were incubated at room temperature for 10 min before being centrifuged at 14,000 × g for 10 min at room temperature to pellet insoluble material. Following centrifugation, the supernatant was carefully collected and mixed with 0.2 volumes of SDS sample buffer composed of 5% SDS, 30% glycerol, 5% 2-mercaptoethanol, 100 μg/mL bromophenol blue, and 500 mM Tris-HCl (pH 6.8).

### Immunoblot Analysis

For immunoblot analysis, 15 μL of each protein extract was separated on either a 10% or 12% SDS–polyacrylamide gel and subsequently transferred onto a PVDF membrane (Bio-Rad, Hercules, CA, USA). Membranes were blocked for 30 min at room temperature in TBS-T (150 mM NaCl, 10 mM Tris-HCl, pH 7.5, and 0.1% Tween-20) supplemented with 5% (w/v) skim milk.

Following blocking, membranes were incubated overnight at 4 °C with primary antibodies diluted in TBS-T containing 5% skim milk. The following primary antibodies were used: rabbit anti-Rh1 (Bentrop et al., 1997), mouse anti-mCherry (Thermo Fisher Scientific, Cat# 740003M), and mouse anti-α-tubulin (DSHB, Cat# E7, RRID:AB_528499; 1:1000 dilution). α-Tubulin was used as the loading control.

After primary antibody incubation, membranes were washed three times for 5 min each with TBS-T and subsequently incubated for 1 h at room temperature with horseradish peroxidase (HRP)-conjugated secondary antibodies diluted 1:10,000 in TBS-T containing 5% skim milk (Sigma-Aldrich, Cat# A9044, RRID: AB_258431). Following three additional washes of 5 min each in TBS-T, immunoreactive signals were visualized using enhanced chemiluminescence (ECL; 0.091 M Tris-HCl, pH 8.6, 0.0227% luminol, and 0.01% H₂O₂). Chemiluminescent signals were acquired with a ChemiDoc XRS+ Molecular Imager and analyzed using Quantity One 4.6.9 software (Bio-Rad).

Quantification of immunoblot signals was performed using Image Lab 4.0 software (Bio-Rad) by integrating the pixel intensities of individual protein bands. The signal intensities of Rh1 and Rh1::eGFP::mCherry were normalized to the corresponding α-tubulin signal from the same sample. The normalized Rh1 signal intensity from wild-type flies was set to 100%. Each data point represents three independent biological replicates. Statistical analyses were performed using GraphPad Prism version 10.0.2 (GraphPad Software, San Diego, CA, USA; RRID: SCR_002798). Differences between experimental groups were evaluated by one-way ANOVA followed by Tukey’s multiple-comparison post hoc test. Statistical significance is indicated in the corresponding graphs by horizontal bars connecting the compared groups and is represented as follows: ns, not significant; **P < 0.002; ***P < 0.0002.

### Electroretinography (ERG)

Electroretinogram (ERG) recordings were performed on flies that had been briefly anesthetized on ice and mounted in custom-made holders prepared from 200 μL pipette tips. All recordings were carried out with the mounted flies positioned centrally inside a Faraday cage to reduce electrical interference.

Recording and reference electrodes were prepared from glass micropipettes filled with Davenport solution (100 mM NaCl, 2 mM KCl, 1 mM CaCl₂, and 1.8 mM NaHCO₃, pH 7.2). Micropipettes were pulled from BF150-75-10 glass capillaries (Sutter Instrument, Tuttlingen, Germany) using a PC-10 micropipette puller (Narishige Scientific Instruments, Tokyo, Japan). Chlorinated silver wires (250 μm diameter; Goodfellow, Huntingdon, UK) were inserted into the micropipettes to serve as recording electrodes. Chlorination of the silver wires was performed with an ACl-01 automatic chlorider (NPI Electronics, Tamm, Germany). The resulting electrodes exhibited a resistance of approximately 30 mΩ when measured in 0.9% NaCl solution.

The recording electrode was positioned beneath the corneal surface of the compound eye, whereas the reference electrode was inserted into the head capsule. ERGs were recorded at room temperature after 3 min of dark adaptation. Light stimulation was delivered using a PLED-02M stimulator (NPI Electronics) connected to either a 590 nm orange LED or a 470 nm blue LED (Roithner Lasertechnik, Vienna, Austria). The light beam was focused through two collimating lenses (Linos, Göttingen, Germany), resulting in a light intensity of 3.58 mW/cm² at the level of the fly eye.

Two stimulation paradigms were employed. The first consisted of the OBBOO protocol, comprising 5 s of orange light, 10 s of darkness, 5 s of blue light, 10 s of darkness, a second 5 s blue-light stimulus, 10 s of darkness, 5 s of orange light, 10 s of darkness, and a final 5 s orange-light stimulus. The second paradigm consisted of a single 5 s orange-light stimulus followed by 30 s of darkness.

Electrical signals were amplified using an EXT 10-2F amplifier (NPI Electronics) equipped with a 700 Hz low-pass filter. Data acquisition and subsequent analysis were carried out with WinWCP version 4.7.6 software (University of Strathclyde, Glasgow, Scotland; RRID:SCR_014713).

### Immunostaining

For cryosectioning, fly heads were separated from the bodies, bisected, and fixed in 2% paraformaldehyde (PFA) prepared in phosphate-buffered saline (PBS; 137 mM NaCl, 2.7 mM KCl, 10 mM Na₂HPO₄, 2 mM KH₂PO₄, pH 7.2) for 30 min at room temperature. Following fixation, samples were washed twice for 5 min each in phosphate buffer (77 mM Na₂HPO₄ and 23 mM NaH₂PO₄, pH 7.4).

Tissue cryoprotection was achieved by sequential sucrose infiltration. Samples were incubated in 10% sucrose for 45 min, followed by 25% sucrose for 30 min at room temperature, and subsequently infiltrated overnight at 4 °C in 50% sucrose prepared in phosphate buffer. The tissues were then embedded in Tissue-Tek® O.C.T. compound (Sakura, Tokyo, Japan) and cryosectioned at −25 °C using a CryoStar NX50 cryostat (Epredia). Sections with a thickness of 10 μm were collected and post-fixed in 2% PFA for 5 min, followed by three washes in PBS for 5 min each.

For immunofluorescence staining, sections were blocked for 2 h at room temperature in PBS-T containing 1% (w/v) bovine serum albumin (BSA) and 0.3% (v/v) Triton X-100. Primary antibodies diluted in PBS-T were applied overnight at 4 °C. The following primary antibodies were used: mouse anti-Rab7 (DSHB(AB_2722471)), mouse anti-TRP (DSHB(MAb83F6)) and rabbit anti-Calnexin (DSHB(AB_2722011)). After incubation with the primary antibodies, sections were washed three times for 5 min each with PBS and subsequently incubated for 2 h at room temperature with Alexa Fluor-conjugated secondary antibodies (Thermo Fisher Scientific, Cat# A-11045 and A-21244) diluted in PBS-T. For conventional fluorescence microscopy, Alexa Fluor 680-conjugated secondary antibodies was used at a dilution of 1:100. Following three additional 5 min washes in PBS, sections were mounted in Mowiol 4-88 supplemented with 2% *n*-propyl gallate as an antifade reagent.

Fluorescence images were acquired using an AxioImager.Z1m microscope (Carl Zeiss, Jena, Germany) equipped with an ApoTome optical sectioning module. Images were recorded using either an EC Plan-Neofluar 40×/1.3 Oil objective in combination with an Axiocam 530 mono camera (Carl Zeiss). Image acquisition was performed using ZEN 2 (Blue Edition) software (Carl Zeiss).

## ACKNOWLEDGEMENTS

The authors are grateful to Armin Huber for initiating this project and for critical reading of the manuscript. They thank Johannes Gibhardt (University of Hoheneheim) for providing the mCherry plasmid used for cloning and the *Drosophila* Injection Service at University of Zurich for generating transgenic fly lines.

## AUTHORS CONTRIBUTIONS

N.F. designed the study, performed experiments, analyzed data, wrote the manuscript and prepared figures; H.G. performed experiments, analyzed data and prepared figures; C.S. performed experiments.

## REFERENCES

1. Alloway PG, Howard L, Dolph PJ. The formation of stable rhodopsin-arrestin complexes induces apoptosis and photoreceptor cell degeneration. Neuron. 2000;28(1):129–138.

2. Bentrop J, Schwab K, Pak WL, Paulsen R. Site-directed mutagenesis of highly conserved amino acids in the first cytoplasmic loop of *Drosophila* Rh1 opsin blocks rhodopsin synthesis in the nascent state. EMBO J. 1997;16:1600–1609.

3. Chinchore Y, Mitra A, Dolph PJ. Accumulation of rhodopsin in late endosomes triggers photoreceptor cell degeneration. PLoS Genet. 2009;5(2):e1000377.

4. Colley NJ, Cassill JA, Baker EK, Zuker CS. Defective intracellular transport is the molecular basis of rhodopsin-dependent dominant retinal degeneration. Proc Natl Acad Sci USA. 1995;92(7):3070–3074.

5. Feizy N, Leuchtenberg SF, Steiner C, Fliegner L, Würtz B, Huber A, et al. In vivo identification of Drosophila rhodopsin interaction partners by biotin proximity labeling. Sci Rep. 2024; 14:1986. doi:10.1038/s41598-024-52041-3.

6. Han J, Reddig K, Li HS. Prolonged Gq activity triggers fly rhodopsin endocytosis and degradation, and reduces photoreceptor sensitivity. EMBO J. 2007;26(24):4966–4973. doi: 10.1038/sj.emboj.7601929.

7. Hardie RC, Raghu P. Visual transduction in Drosophila. Nature. 2001;413(6852):186–193.

8. Huber A, Wolfrum U, Paulsen R. Opsin maturation and targeting to rhabdomeral photoreceptor membranes requires the retinal chromophore. Eur J Cell Biol. 1994;63(2):219– 229.

9. Iwanami N, Nakamura Y, Satoh T, Liu Z, Satoh AK. Rab6 Is Required for Multiple Apical Transport Pathways but Not the Basolateral Transport Pathway in Drosophila Photoreceptors. PLoS Genetics. 2016;12(2):e1005828.

10. Kimura S, Noda T, Yoshimori T. Dissection of the autophagosome maturation process by a novel reporter protein, tandem fluorescent-tagged LC3. Autophagy. 2007;3(5):452–460.

11. Kiselev A, Socolich M, Vinós J, Hardy RW, Zuker CS, Ranganathan R. A molecular pathway for light-dependent photoreceptor apoptosis in Drosophila. Neuron. 2000;28(1):139–152. doi:10.1016/S0896-6273(00)00092-1.

12. Montell C. Visual transduction in Drosophila. Annu Rev Cell Dev Biol. 1999;15:231–268.

13. O’Tousa JE, Baehr W, Martin RL, Hirsh J, Pak WL, Applebury ML. The Drosophila ninaE gene encodes an opsin. Cell. 1985;40(4):839–850.

14. Ozaki K, Nagatani H, Ozaki M, Tokunaga F. Maturation of major Drosophila rhodopsin, ninaE, requires chromophore 3-hydroxyretinal. Neuron. 1993;10(6):1113–1119.

15. Parenti N, Del Grosso A, Antoni C, Cecchini M, Corradetti R, Pavone FS, Calamai M. Direct imaging of APP proteolysis in living cells. PeerJ. 2017;5:e3086.

16. Satoh AK, Ready DF. Arrestin1 mediates light-dependent rhodopsin endocytosis and cell survival. Curr Biol. 2005;15(19):1722–1733.

17. Satoh T, Nakamura Y, Satoh AK. The roles of Syx5 in Golgi morphology and Rhodopsin transport in *Drosophila* photoreceptors. Biology Open. 2016;5(10):1420–1430. doi:10.1242/bio.020958.

18. Richter D, Katz B, Oberacker T, Tzarfaty V, Belusic G, Minke B, Huber A. Translocation of the Drosophila transient receptor potential-like (TRPL) channel requires both the N- and C-terminal regions together with sustained Ca2+ entry. J Biol Chem. 2011;286:34234–34243. doi:10.1074/jbc.M111.278564.

19. Wang S, Tan KL, Agosto MA, Xiong B, Yamamoto S, Sandoval H, Jaiswal M, Bayat V, Zhang K, Charng WL, David G, Duraine L, Venkatachalam K, Wensel TG, Bellen HJ. The retromer complex is required for rhodopsin recycling and its loss leads to photoreceptor degeneration. PLoS Biol. 2014;12(4):e1001847.

20. Xiong B, Bellen HJ. Rhodopsin homeostasis and retinal degeneration: lessons from the fly. Trends Neurosci. 2013;36(11):652–660. doi:10.1016/j.tins.2013.08.003.

21. Zhao N, Li N, Wang T. PERK prevents rhodopsin degradation during retinitis pigmentosa by inhibiting IRE1-induced autophagy. J Cell Biol. 2023;222(5):e202208147. doi:10.1083/jcb.202208147.

22. Zuker CS, Cowman AF, Rubin GM. Isolation and structure of a rhodopsin gene from Drosophila melanogaster. Cell. 1985;40(4):851–858.

